# Isolation and antibiotic profile of *Vibrio* spp. in final effluents of two wastewater treatment plants in the Eastern Cape of South Africa

**DOI:** 10.1101/330456

**Authors:** Olayinka Osuolale, Anthony Okoh

## Abstract

**Background:** Poorly or partially treated wastewater disposed of can contaminate water and even properly treated sewage can have its problems. The highlight of this danger is wastewater treatment plants serving as reservoir for proliferation of antibiotic resistant organisms. We have reported the state of two wastewater treatment in the Eastern Cape of South Africa which discharge poorly and partially treated effluents. Our aims to identify Vibrio spp. and their antibiotic profiles in treated final effluent discharge from wastewater treatment plant.

**Methods:** Culture based approach using the TCBS agar for isolation *Vibrio* spp., presumptive isolates were purified and confirmed using PCR. The confirmed isolated were also genotyped to identify the species present. The antibiotic profiling of the confirmed isolates was using the CLSI recommended first line antibiotics for Vibrio.

**Results:** Out of the 786 presumptive isolates, 374 were confirmed as *Vibrio* spp. None of the Vibrio spp. pathotypes were present in the confirmed isolates. Randomized isolates of 100 Vibrio spp. were selected, > 90 % of the isolates were susceptible to Ciprofloxacin, and > 50 – 80 % for Ampicillin, Chloramphenicol, Tetracycline, Cefotaxime, and Trimethoprim-sulfamethoxazole respectively.

**Conclusions:** We are able to isolate Vibrio spp. from treated effluents but none of their pathotypes were present. The antibiotic agents considered for primary testing which are ciprofloxacin was the most effective of the antibiotic drugs, followed by cefotaxime, tetracycline with less susceptibility. Contamination from discharged effluents from wastewater treatment can lead to spread of spread of disease in this environment. The WWTPs studied are sources of pollution to surface water with environmental and public health.

## INTRODUCTION

Vibrio are gram negative, rods and are motile with a polar flagellum containing diverse groups [1,2]. Members of various species are known to cause acute gastroenteritis infections [3,4], wound infections and primary septicemia [5]. Many *Vibrio* spp. are pathogenic to humans and have been implicated in food-borne disease [6]. They are naturally found in the estuarine and marine environment [7,8]. The aquatic environment have been identified to be a medium of transmission of this organism through which the isolation of the organism has been found in seafood [7,9,10]. The isolation of the microorganism from raw sewerage as well as the final treated effluent showed that wastewater treatment plants do not remove or inactivate all pathogenic microorganisms [11,12] and hence Wastewater has been implicated in the distribution of Vibrio in the environment and surface water [13,14]. Study have shown the presence of high level antibiotic resistance *Vibrio cholera*e in the final effluent of stabilization pond revealing the imminent danger of poorly treated effluent [15]. One of the recent deadliest outbreaks of the organism was in Haiti which was attributed to poor wastewater management and the existence of poor sanitary condition in the country [16,17]. South Sudan and Kenya are recently ravaged by cholera outbreak, and the world worst’s outbreak is in Yemen [18–21]. In South Africa, Vibrio has been isolated from faeces of domestic animals in rural area of Limpopo [22] and in 2002, the province recorded one of its first outbreak of the diseases [23,24]. Between 2002 and 2004, there have been reported cases of the disease outbreak in the Eastern Cape of South Africa [25]. More studies from the region found *Vibrio* spp. isolated from treated effluents of wastewater treatment plants [12,26].

Worldwide, cases of most bacterial pathogens becoming more resistant to commonly used antimicrobial agents are increasing [27]. In developing countries, increase in antimicrobial resistance in enteric pathogens is especially important where diarrhea is common [28]. Multi drug resistance has been reported in effluent from Wastewater treatment in Eastern Cape, which are considered reservoir for antibiotic resistance bacteria [29,30]. Effluent water is still daily discharged into surface water in the Eastern Cape and therefore the essence of this monitoring study is on the antibiotic resistance prevalence of the organism in the final effluent of wastewater treatment plant. This formed part of a large project study done and reported [31–33]. This study build upon the state of knowledge on effluent quality discharge in the Eastern Cape. This study profile approach centered on antimicrobial agents for use in treatment of cholera as recommended by the WHO.

## MATERIALS AND METHODS

The detailed sampling sites for 2 wastewater treatment plants WWTP-A and WWTP-B, sampling collection and processing is as reported and published in Osuolale & Okoh, [32].

### Isolation of Vibrio

Enumerations of presumptive Vibrio pathogens were carried out by the method using sterile TCBS agar plants as described by Bopp et al., [34]. Bacteriological analysis of the effluent samples for bacteria counts and isolation was determined by membrane filtration (47mm, 0.45mm pore size), according to SABS, [35]. Serial dilutions of the samples were prepared. Sample dilutions were homogenate before filtering (100ml). On certain occasions where there was excessive chlorine dosage in the effluent, the raw samples were filtered. The filtered samples were placed on selective agar for the target organisms in triplicates. The plates were allowed 15 minutes to dry, invert, and incubate promptly for 24hrs at 37 °C. After 48 hours incubation, colonies appearing as greenish or yellowish in colour were counted and reported as CFU/100 ml SABS, [35] in suitable range (0-300 colonies). Presumptive Vibrio bacteria isolated from the plates were purified and subjected to molecular identification. Polymerase chain reaction (PCR) was used to confirm the identities of the Vibrio species using the species specific primers as described by Tarr et al., [1].

### Phenotypic identification of Vibrio

Considering the salt tolerance of some *Vibrio* spp., can grow at a salt concentration of 3% NaCl [36]. Presumptive isolates from the TCBS culture plates were inoculated into a tube each of 1% tryptone broth (TSB) with 2% NaCl and incubated 18-24 h at 35-37 °C. Profuse growths in tubes are considered as positive. Various species have different salt tolerance that can be used for identification. This test helps to eliminate presumptive colonies from the TCBS plate which resemble Vibrio, e.g. Proteus [37,38].

### Isolation of genomic DNA and Genotypic identification of Vibrio

Vibrio isolates from the freeze storage were inoculated on TSB broth overnight for crude DNA extraction. Frozen cells were kept on ice to reduce thawing by scraping the ice surface with a loop. ZR Fungal/Bacterial DNA MiniPrep by Zymo Research was used to extract genomic DNA following the manufacturer’s instruction. The genomic extracts were immediately used in the molecular identification of the isolated organisms. Primers specific for the confirmation of the Vibrio isolates was used in the polymerase chain reaction. PCR amplification was performed with a MyCycler thermal cycler PCR (Bio-Rad). The PCR solution contained 2 x PCR mastermix, 100uM each of 0.2 to 0.5 uM each of the primers. The total volume for PCR reaction was 25 µl, 5 µg of template DNA from each bacterial strain was added to make the final 25ul reaction volume. PCR confirmation reactions were performed to amplify the 16sRNA IGS regions of *Vibrio* spp. by using V. 16S-700F (CGG TGA AAT GCG TAG AGA T) and V. 16S-1325R (TTA CTA GCG ATT CCG AGT TC) primers of 663 bp. Confirmed isolates were further subjected to genotypic identification for *V. parahaemolyticus, V. vulnificus* and *V. fluvialis*. The positive control was from Leibniz-Institut DSMZ (GmBH). The cycling conditions was: a 15 mins initial denaturation at 93 °C followed by 35 cycles of 92 °C for 40 sec, 57 °C for 1 min, and 72 °C for 1.5 mins and a final soak at 72 °C for 7 mins [1]. Gel electrophoresis was performed on the PCR products and ran on a 2% w/v agarose gel at 100 V for approximately 90 mins. Gel images was captured digitally and analyzed using the Uvitec, Alliance 4.7.

### Antimicrobial susceptibility

The antibiotic susceptibility testing for Vibrio isolates was determined using the following antibiotic discs: ampicillin (10 µg), tetracycline (30 µg), chloramphenicol (30 µg), cefotaxime (30 µg), Trimethoprim-sulfamethoxazole (1.25/23.75 µg), and ciprofloxacin (5 µg). The choice of antibiotics was based on recommended drug for primary testing of Vibrio spp. by CLSI [40,41].

## RESULTS

The outcome of the PCR analysis was able to augment the culture-based method employed in the detection of the Vibrio isolates. The confirmation of the target gene of interest on the presumptive isolates validated the presence of Vibrio in the final effluents of the wastewater treatment plants studied. Some of the tested samples showed consistency with the expected band size of 663bp on the ladder and a positive control as a guide (Fig. 1).

**Figure 1.**
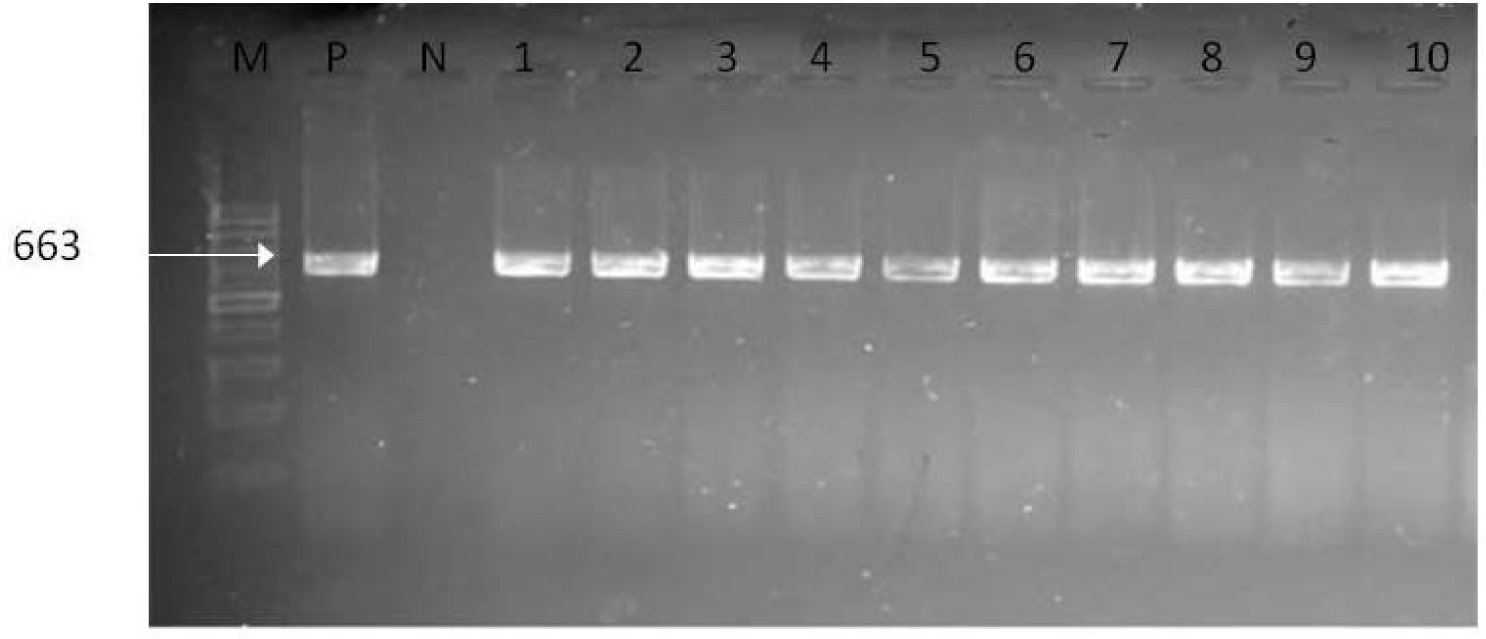
Agarose gel electrophoresis of 16s rRNA gene amplification products of Vibrio. M: Molecular weight marker (100bp) P: Positive control N: Negative control; Lanes 1-10: *Vibrio* spp. isolates

The target gene 16S rRNA was positive for some of the presumptive isolates. 207 out of 340 isolates from WWTP-A site were positive for these the gene, while 167 out of the 446 isolates from WWPT-B were confirmed for the 16S rRNA gene.

A total of 100 confirmed PCR isolates were selected for used in investigation for their antibiotic resistance. The antibiotic resistance profile test was done using the disc diffusion susceptibility testing and the zones of inhibition was compared against the CLSI standard [40,41]. The final interpretation of measurements, grouped into three categories, namely sensitive, intermediate and resistance, is summarized in Table 2.

The specific readings of inhibition zone diameter, the intrinsic resistance or susceptibility of each testing isolates, were found to differ even to the same antibiotics, this variable susceptibility/resistance was observed in all the antibiotics tested. For instance, although all 100 isolates were interpreted as either resistance or sensitive to tetracycline, isolates that are susceptible to tetracycline are as well considered susceptible to doxycycline and minocycline. However, some isolates that are intermediate or resistant to tetracycline may be susceptible to, minocycline, or both.

The quality control steps were as recommended by [40,41]. The tested isolates were susceptible to Ciprofloxacin (92%), Trimethoprim-sulfamethoxazole (80%), Cefotaxime (79%), Chloramphenicol (67%), Ampicillin (58%) and Tetracycline (54%) with the least susceptibility. In addition, 71.6% of all the isolates tested remained susceptible to all antimicrobials, 6.8%% with intermediate susceptibility and 21.5% exhibited resistance to all the antimicrobials.

## DISCUSSION

In this study, the WWTP-A treatment plant had a high prevalence of Vibrio observed and in contrast to WWTP-B which had a very low prevalence of the organism in the sample analyzed. The data is not shown here but published in one of our article [32]. An existing report on WWTP-B by Igbinosa [12], also isolated *Vibrio* spp. from the final effluent of the wastewater treatment plant. In another area of South Africa in Gauteng, Vibrio was found in the final effluent of the wastewater plant [42]. In a work done by Ye & Zhang, [43] in Hong Kong, also found high prevalence of Vibrio in the effluent of the studied treatment plant. Evaluating the treatment technologies used show that the activated sludge system was far more effective in reducing the Vibrio pathogen than the biofilter/trickling filter system. The results coincide with the report of Ngari, Kotut, & Okemo, [44] which had effluent from trickling filter having low removal rate of pathogens. In contrast, Ramteke et al., [45] found the activating sludge system to have high removal rate of Vibrio. High level of chlorination (high free chlorine) was observed for some periods during samplings in the both site’s plants. High free chlorine was more frequent in WWTP-B wastewater treatment plant than in WWTP-A wastewater plants (data not shown). Vibrio was persistently isolated from the high chlorinated effluent in WWTP-B though at low concentration less than it was at WWTP-A. The frequency of the Vibrio isolation was observed more at the WWTP-B discharge point than at the final effluent point. Even at the recommended free chlorine level, Vibrio was isolated. This trend was observed in the final effluents in some of wastewater treatment plants in the Eastern Cape which included studying other pathogens apart from Vibrio [3,12,26,46] and factors such as contact time, temperature, pH may affect the efficiency of the disinfectants [47]. The presence of organic compounds and ammonia in the effluent also contribute to the ineffectiveness of the disinfection process [48,49]. It is therefore important that the effluent be of high quality for maximum effect of the disinfectant [50].

Coupled with the under performance of the WWPT-A treatment plant in eliminating the pathogen, organisms are being re-introduced back into the environment and this can create a vicious cycle of outbreak of infections from the infectious organisms. The Green Drop status, which implies excellent wastewater management and a respect for the environment and the health of the community at large, is given to municipalities that comply with good wastewater discharge standards for 90% of the time [51]. The previous Green Drop status reports of WWPT-A treatment plant was awarded a medium risk plant between 2010-11 and 2012 [52,53] and the most recent reports showed no changed in their treatment processes for 2013 and 2014 [54,55]. The outcome of this current study on the plant showed it is a high risk plant with potential danger to the environment, therefore, the plant needs urgent attention. The Green Drop of 2012 also identified some of the challenges facing the WWPT-A Plant which included effluent non-compliance and operating capacity that exceeds design capacity [53]. In contrast, the green status for WW-Dim went from a medium risk rating to a low risk rating [52,53]. The effluent quality of the plant also showed it as a low risk wastewater plant. However, the detection of *Vibrio* spp., though at a very minimal level, is of concern judging from the nature of the organism as one having the potential to initiate epidemic infection.

The samples positive for *Vibrio* spp. were further screened for the *V. parahaemolyticus, V. vulnificus and V. fluvialis* pathotypes. All the screened isolates were negative for the tested Vibrio pathotypes. The target genes (Table 1) specific for the identification of these pathotypes were not detected in the tested isolates. With the exception of V. cholera which could not be tested the tested strains are not ubiquitous to the natural fresh or salt aquatic environment as are the V. cholera [11]. *V. fluvialis, V. parahaemolyticus and V. vulnificus* are reported as the most frequently encountered pathogenic Vibrios in marine environments, coastal, estuaries and brackish waters as well as seafood, which is considered a natural habitat for this strains of *Vibrio* spp. [4,56]. In contrast to our work is the study done by Igbinosa, [12] who reported the presence of the *V. fluvialis, V. parahaemolyticus and V. vulnificus* in the final effluent of a wastewater treatment plant. The prevalence of these pathogenic Vibrio in the environment were reported to be influenced by temperature and salinity and the concentration of salinity differs for each *Vibrio* spp. at which they can survive [5,9]. The public health consequences of these pathogenic organisms cannot be over emphasized as all these strains have been attributed to human diseases [1] and are known to cause gastrointestinal disease syndrome [57].

**Table 1.**
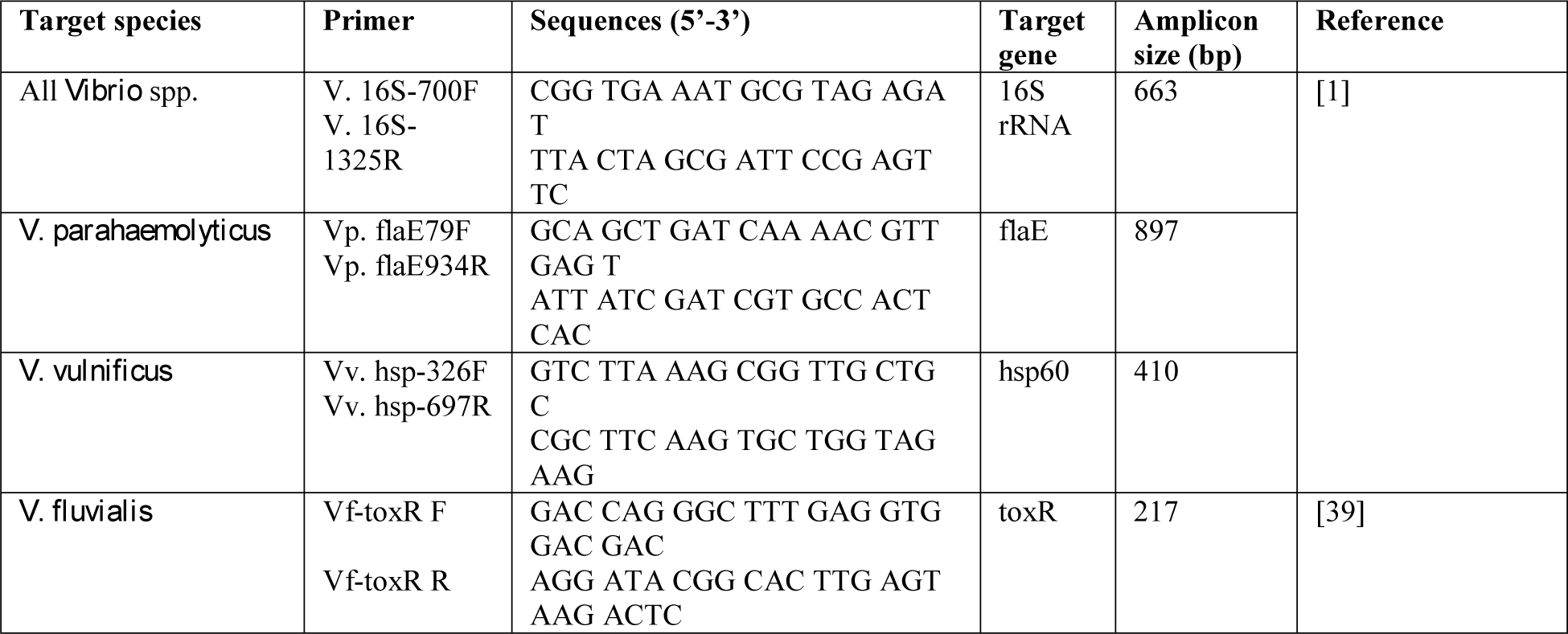
Primer pairs for Vibrio confirmation, genotypes, and expected amplicon size

**Table 2.** Susceptibility profile of randomly selected 100 Vibrio isolates to various antibiotics.

| Antimicrobial agents | Number (%) of isolates = 100 |  |  |
| --- | --- | --- | --- |
|  | Susceptible | Intermediate | Resistant |
| Ampicillin (10 µg) | 58 | 13 | 29 |
| Chloramphenicol (30 µg) | 67 | 14 | 19 |
| Ciprofloxacin (5 µg) | 92 | 4 | 4 |
| Tetracycline (30 µg) | 54 | 3 | 43 |
| Cefotaxime (30 µg) | 79 | 4 | 17 |
| Trimethoprim-sulfamethoxazole (1.25/23.75 µg) | 80 | 3 | 17 |

### Antimicrobial susceptibility testing for Vibrio spp

*Vibrio* spp. are considered to be significant infectious pathogens which are the causative agents for vibrosis [58,59]. They are characterized by diarrhea, wound infections, primary septicemia, and gastroenteritis or other extra-intestinal infections related to exposure to contaminated sources [60]. Most isolates tested in this study were susceptible to the antimicrobial agents recommended for primary testing by CLSI [40]. Treatment recommendations for Vibrio infections include: tetracycline (doxycycline, tetracycline), fluoroquinolones (ciprofloxacin, levofloxacin), third-generation cephalosporins (cefotaxime, ceftazidime, ceftriaxone), aminoglycosides (amikacin, apramycin, gentamicin, streptomycin) and folate pathway inhibitors (trimethoprim-sulfamethoxazole) [61,62]. All *Vibrio* spp. studied here showed some degree of resistance to all the antibiotics used for testing. In the present study, data on antibiotic resistant zones indicate that all the 100 isolates of *Vibrio* spp. were 38% resistant to tetracycline, 26% to ampicillin, 16% to chloramphenicol, 14% to cefotaxime, 13% to trimethoprim-sulfamethoxazole and 1% to ciprofloxacin.

The result of antimicrobial susceptibility testing showed Vibrio isolates had the highest susceptibility of 92% to ciprofloxacin. Similar susceptibility level to ciprofloxacin was reported by Ismail et al., [63] during a study of *Vibrio cholera* outbreak in South Africa. About 90% of Vibrio isolated from river water used as water sources in a rural communities of Venda in South Africa were susceptible to ciprofloxacin [64]. Igbinosa, [12] worked on *Vibrio* spp. isolated from the final effluent of a wastewater treatment plant in South Africa reported susceptibility in the range of 70% to 90% to the antibiotics. Report from other studied areas show similar susceptibility level at 100% susceptible [62,65–67] as compared to the 92% recorded in this study while 96.4% susceptibility was reported by Benedicta, [68]. Other *Vibrio* spp. which is mostly associated sea animals and marine environments are also reported to be susceptible to the antibiotic [69]. Reduced susceptible has been reported in two West Africa countries [70,71], India [72] and Bangladesh [73] showing the possibility of the organism to develop resistance. It was highlighted in the Leclercq et al., [74], when antibiogram with quinolones are read, resistance to the most active fluoroquinolone in vitro indicates resistance to all fluoroquinolones in both Gram-negative and Gram-positive organisms An exception to this rule in Gram-negative organisms is the potential production of the AAC(6’)-Ib-cr enzyme, which affects ciprofloxacin but not levofloxacin [74]. Kim et al., [75] were able to demonstrate the transferrable of quinolone resistance gene in *Vibrio cholera* and thus highlights the opportunities for gene exchange among bacteria living in aquatic environments which can confer resistance.

Trimethoprim-sulfamethoxazole or co-trimoxazole has an appreciable level of susceptibility of about 80%. Report by Shaw et al., [62] showed full susceptibility to trimethoprim-sulfamethoxazole. This is however in sharp contrast to the susceptibility of Vibrio to the antibiotic previously reported where high level of resistance has been documented for Vibrio isolates from South Africa [63,76,77], India [72,78] and the Vietnam [67]. Likewise multiple resistances has been reported for antibiotics tested along with co-trimoxazole which include ampicillin [72,77,79]. This as well corroborate with the work done by Igbinosa, [12] who reported ampicillin resistance to *Vibrio* spp. isolated from the final effluent at the Eastern Cape, South Africa. This study recorded 58% susceptibility level with 29% resistance to ampicillin. This compared favorably with other studies which exhibiting resistance to ampicillin [61,80,81]. Furthermore, Igbinosa, [12] and Quilici et al., [71] reported intermediate susceptibility to ampicillin while Ismail et al., [63] and Tran et al., [67] reported high susceptibility level to ampicillin from *Vibrio cholera* tested in South Africa and Vietnam. Ismail et al., [63] was conversely quick to point out the observed change to the susceptibility of some of the same isolate tested on ampicillin to have suddenly become resistance though at a very minimal resistance level, further showing the rapid development antibiotics resistance of *Vibrio cholera* in South Africa. In addition, the CLSI publication stated result for ampicillin can be used to predict for amoxicillin [82]. It can therefore be deduced that amoxicillin will as well have variant susceptibility or resistance to *Vibrio* spp. Susceptibilities of Vibrio spp. have been shown to vary by species, particularly with regard to the older penicillin, cephalosporins, and sulfonamides [83].

In contrast to our study, we had an susceptibility of 54% to tetracycline while [71] reported high susceptibility to the antibiotic. There were others who reported intermediate resistant to tetracycline [67], and co-multiple resistances to tetracycline and chloramphenicol [76–79]. According to the CLSI documentation, Organisms susceptible to Tetracycline are also considered susceptible to doxycycline and minocycline. However, some organisms that are intermediate or resistance to tetracycline may be susceptible to doxycycline or minocycline or both [82]. Tran et al., [67] are able to show in their work susceptibility to doxycycline by *Vibrio cholera*e with intermediate resistance.

The observed susceptibility level for chloramphenicol was 64% as against 100% susceptibility recorded from other others [63,67,71] and Igbinosa, [12] reported 100% resistance to the chloramphenicol. *Vibrio* spp. across different studies showed variants levels of susceptibility to resistance to chloramphenicol [84,85], and the presence of chloramphenicol resistance gene was identified in Vibrio isolates showing resistance to the drug [86].

Cefotaxime is a third generation cephalosporin and are still largely effective against *Vibrio* spp. [87]. There have been reports of *Vibrio* spp. resistance to cefotaxime though at a very minimal level [88]. The susceptibility to cefotaxime observed in our study is high but also with 17% resistant isolates. Liang et al., [89] reported a single isolate resistance to cefotaxime while three isolates showed intermediate reactions. Similarly, Shaw et al., [62] also reported intermediate resistance to cefotaxime in their study. Most studies have shown high susceptibility of *Vibrio* spp. to cefotaxime as reported in the work done by Han et al., [61] and Zanetti et al., [90]. The use of cefotaxime with minocycline for treatment of some of *Vibrio* spp. like Vibrio vulnificus is recommended for effectiveness and were found to act synergistically in inhibiting the organism [91].

Wastewater treatment plant becoming source of antibiotic reservoir for bacteria is a concern. The increasing threat posed by them stem from mismanagement of the treatment processes. Treated effluent studied by Larsson, de Pedro, & Paxeus, [92] found that effluent does constitute major environmental problem as a result pharmaceutical products from industrial wastes. Li et al., [93] report the presence of Penicillin G and its degraded products in effluent sample analysed. The presence of Fluoroquinolones and sulfamethoxazole have also been detected in the effluent of some wastewater plants in the western cape of South Africa [94]. However it is yet to be agreed upon if wastewater treatments plants is an important source in the emergence of resistant bacteria in the environment, i.e. is the concentration of the antibiotic and the bacterial density high enough, is the exposure long enough to promote resistance or to select resistant bacteria [95]. Though it has been shown that Bacteria can acquire multidrug resistance through sequential transfer of multiple-resistance determinants located on mobile genetic elements [96] their ability to do so in wastewater treatment plants are yet to be fully established [95]. Though recent report by Luo et al., [97] confirmed the possibility of gene transfer from resistant organism to indigenous organisms in wastewater treatment plant. Ohlsen et al., [98] indicate that the transfer of resistance and the selection of resistant bacteria are not favored at antibiotic concentrations as high as those found in hospital effluents or the aquatic environment. Most resistance organisms found in the environment are believed to have come from previously resistant organism. In the review work of Kümmerer, [95] showed that resistant organisms were both isolated from effluents which received hospital waste and municipal influent. This debunks the view that resistant organisms would have been more prevalent in plant receiving hospital waste as compared to municipal plant. Some of the resistant isolates observed in this study could as well as arise from the surrounding communities since there is no source of pharmaceutical industrial influents or hospital waste into the plants. The fallback will be on the use of antibiotic within the communities as this has been reported as source of antibiotics in the environment [99] and the occurrence of antibiotics may promote the selection of antibiotic resistance genes (ARGs) and antibiotic resistant bacteria (ARB), which pose health risks to the environment, humans and animals [100]. Dalsgaard et al., [76] demonstrated multiple-drug resistant V. cholerae o1 isolates showing resistance to all the antibiotics traditionally used to treat cholera which is disturbing and has a direct impact on the treatment of current and future cholera cases in South Africa and other countries to which this isolate may spread.

Therefore, continued monitoring of both the prevalence and the antimicrobial susceptibility profile is important to better ensure environmental safety; particularly single resistance to ciprofloxacin observed against Vibrio also limit treatment effectiveness and should be monitored. As most of the antimicrobial agents recommended for treatment of E. coli and Vibrio illnesses by CLSI showed some form of resistances is likely to be problematic. Based on our data, treatment of illnesses may benefit from the use of meropenem that was 100% effective against E. coli and ciprofloxacin which was the only antibiotics that was 99% effective against *Vibrio* spp. in this study.

## AVAILABILITY OF DATA AND MATERIALS

The data that support the findings of this study are available from Water Research Commission (WRC), South Africa, but restrictions apply to the availability of these data, which were used under license for the current study, and so are not publicly available. Data are however available from the authors upon reasonable request and with permission of WRC. As the data originate from a clinical data set it was not possible to obtain consent for publication of individual patient data.

## AUTHORS’ CONTRIBUTIONS

O. O made substantial contributions to acquisition of data analyzed the data, wrote the manuscript. A. I designed and supervised the study. All authors read and approved the final manuscript and agreed to be accountable for all aspects of the work in ensuring that questions related to the accuracy or integrity of any part of the work are appropriately investigated and resolved.

## REFERENCE

1. Tarr CL, Patel JS, Puhr ND, Sowers EG, Bopp CA, Strockbine NA. Identification of Vibrio isolates by a multiplex PCR assay and rpoB sequence determination. J. Clin. Microbiol. 2007;45:134–40.

2. Noorlis A, Ghazali F, Cheah Y. Prevalence and quantification of Vibrio species and Vibrio parahaemolyticus in freshwater fish at hypermarket level. Int. Food Res 2011;695:689–95.

3. Igbinosa EO, Obi LC, Okoh AI. Occurrence of potentially pathogenic vibrios in final effluents of a wastewater treatment facility in a rural community of the Eastern Cape Province of South Africa. Res. Microbiol. Elsevier Masson SAS; 2009;160:531–7.

4. Ramamurthy T, Chowdhury G, Pazhani GP, Shinoda S. Vibrio fluvialis: an emerging human pathogen. Front. Microbiol. Frontiers; 2014;5:91.

5. Amin R, Salem A. Specific detection of pathogenic Vibrio species in shellfish by using multiplex polymerase chain reaction. Glob. Vet. 2012;8:525–31.

6. Peterkin PI. Compendium of methods for the microbiological examination of foods (3rd edn). Trends Food Sci. Technol. 1993;4:199.

7. Johnson CN, Flowers AR, Noriea NF, Zimmerman AM, Bowers JC, DePaola A, et al. Relationships between environmental factors and pathogenic Vibrios in the Northern Gulf of Mexico. Appl. Environ. Microbiol. 2010;76:7076–84.

8. WHO WHO. WHO | Guidelines for drinking-water quality, fourth edition. Geneva: World Health Organization; 2011.

9. Maugeri TL, Caccamo D, Gugliandolo C. Potentially pathogenic vibrios in brackish waters and mussels. J. Appl. Microbiol. 2000;89:261–6.

10. Adeleye I, Daniels F, Enyinnia V. Characterization and pathogenicity of Vibrio spp. contaminating seafoods in Lagos, Nigeria. Internet J. Food Saf. 2010;12:1–9.

11. Eddabra R, Moussaoui W, Prévost G, Delalande F, Dorsselaer A, Meunier O, et al. Occurrence of Vibrio cholerae non-O1 in three wastewater treatment plants in Agadir (Morocco). World J. Microbiol. Biotechnol. 2010;27:1099–108.

12. Igbinosa E. Surveillance of invasive vibrio species in discharged aqueous effluents of wastewater treatment plants in the Eastern Cape province of South Africa. University of Fort Hare; 2010.

13. Curtis T. The fate of Vibrio cholerae in wastewater treatment systems. Cholera Ecol. Vibrio cholerae. Dordrecht: Springer Netherlands; 1996. p. 295–332.

14. Cañigral I, Moreno Y, Alonso JL, González A, Ferrús MA, Ferrús M a. Detection of Vibrio vulnificus in seafood, seawater and wastewater samples from a Mediterranean coastal area. Microbiol. Res. 2010;165:657–64.

15. Mezrioui N, Oufdou K. Abundance and antibiotic resistance of non-O1 Vibrio cholerae strains in domestic wastewater before and after treatment in stabilization ponds in an arid region (Marrakesh, Morocco). FEMS Microbiol. Ecol. The Oxford University Press; 1996;21:277–84.

16. Harris J, LaRocque R, Qadri F, Ryan E, Calderwood S. Cholera. Lancet. 2012;379:2466–76.

17. Michael S. Clean Water, Sanitation Ease Cholera in Haiti Medpage Today [Internet]. MEDPAGETODAY. 2013 [cited 2017 Jul 6]. Available from: http://www.medpagetoday.com/InfectiousDisease/GeneralInfectiousDisease/42265

18. BBC. Kenya cholera outbreak hits dozens at health conference - BBC News [Internet]. BBC. 2017 [cited 2017 Jul 6]. Available from: http://www.bbc.com/news/world-africa-40369318

19. Colin D. Yemen Now Faces “The Worst Cholera Outbreak In The World,” U.N. Saysl]: The Two-Wayl]: NPR [Internet]. NPR. 2017 [cited 2017 Jul 6]. Available from: http://www.npr.org/sections/thetwo-way/2017/06/24/534236954/yemen-now-faces-the-worst-cholera-outbreak-in-the-world-u-n-says

20. Delling; Kadugli; Ed D. Cholera in South Kordofan: Student dies, infections spread Radio Dabanga [Internet]. DABANGA. 2017 [cited 2017 Jul 6]. Available from: https://www.dabangasudan.org/en/all-news/article/cholera-in-south-kordofan-student-dies-infections-spread

21. Al-Mekhlafi HM. Yemen in a Time of Cholera: Current Situation and Challenges. Am. J. Trop. Med. Hyg. 2018;2016:1–8.

22. Keshav V, Potgieter N. Detection of Vibrio cholerae O1 in animal stools collected in rural areas of the Limpopo Province. 2010;36:167–71.

23. Chabala H. A Report on Cholera Outbreak Response in Limpopo Province, Cholera in the Limpopo Province, Epidemic preparedness and outbreak response, Emerging and Re-emerging Infectious Diseases, Communicable Disease Control, National Department of Health. Republic of [Internet]. 2002 [cited 2017 Jul 13]. Available from: https://web.archive.org/web/20130511111654/www.doh.gov.za/show.php?id=1608

24. Anon. Cholera outbreak in Limpopo Province in 2002 Health24 [Internet]. health24. 2007 [cited 2017 Jul 6]. Available from: http://www.health24.com/Medical/Cholera/Learn-from-previous-outbreaks/Cholera-outbreak-in-Limpopo-Province-in-2002-20120721

25. Lehohla P. Provincial Profile 2004 Eastern Cape. Stat. South Africa. Pretoria Stat. South Africa. 2006.

26. Momba M, Osode A, Sibewu M. The impact of inadequate wastewater treatment on the receiving water bodies – Case study: Buffalo City and Nkokonbe Municipalities of the Eastern Cape Province. Water SA. 2009;32:687–92.

27. Roca I, Akova M, Baquero F, Carlet J, Cavaleri M, Coenen S, et al. The global threat of antimicrobial resistance: Science for intervention. New Microbes New Infect. 2015. p. 22–9.

28. Jafari F, Hamidian M, Rezadehbashi M, Doyle M, Salmanzadeh-Ahrabi S, Derakhshan F, et al. Prevalence and antimicrobial resistance of diarrheagenic Escherichia coli and Shigella species associated with acute diarrhea in Tehran, Iran. Can. J. Infect. Dis. Med. Microbiol. 2009;20:e56–62.

29. Okoh AI, Igbinosa EO. Antibiotic susceptibility profiles of some Vibrio strains isolated from wastewater final effluents in a rural community of the Eastern Cape Province of South Africa. BMC Microbiol. 2010;10:143.

30. Igbinosa EO, Obi LC, Tom M, Okoh AI. Detection of potential risk of wastewater effluents for transmission of antibiotic resistance from Vibrio species as a reservoir in a peri-urban community in South Africa. Int. J. Environ. Health Res. 2011;21:402–14.

31. Osuolale O, Okoh A. Human enteric bacteria and viruses in five wastewater treatment plants in the Eastern Cape, South Africa. J. Infect. Public Health. 2017;

32. Osuolale O, Okoh A. Assessment of the physicochemical qualities and prevalence of Escherichia coli and vibrios in the final effluents of two wastewater treatment plants in South Africa: Ecological and public health implications. Int. J. Environ. Res. Public Health. Multidisciplinary Digital Publishing Institute; 2015;12:13399–412.

33. Osuolale O, Okoh A. Incidence of human adenoviruses and Hepatitis A virus in the final effluent of selected wastewater treatment plants in Eastern Cape Province, South Africa. Virol. J. Virology Journal; 2015;12:98.

34. Bopp C, Ries A, Wells J. Laboratory methods for the diagnosis of epidemic dysentery and cholera. 1999;

35. SABS. SANS 5221l]: 2011 SOUTH AFRICAN NATIONAL STANDARD Microbiological analysis of water — General test methods. 2011;7–11.

36. Kaysner C, DePaola A. Vibrio. Bacteriol. Anal. Man. FDA: Center for Food Safety and Applied Nutrition; 2004.

37. McCormack WM, DeWitt WE, Bailey PE, Morris GK, Soeharjono P, Gangarosa EJ. Evaluation of thiosulfate citrate bile salts sucrose agar, a selective medium for the isolation of Vibrio cholerae and other pathogenic vibrios. J. Infect. Dis. 1974;129:497–500.

38. Lotz MJ, Tamplin ML, Rodrick GE. Thiosulfate-citrate-bile salts-sucrose agar and its selectivity for clinical and marine vibrio organisms. Ann. Clin. Lab. Sci. 1983;13:45–8.

39. Chakraborty R, Sinha S, Mukhopadhyay AK, Asakura M, Yamasaki S, Bhattacharya SK, et al. Species-specific identification of Vibrio fluvialis by PCR targeted to the conserved transcriptional activation and variable membrane tether regions of the toxR gene. J. Med. Microbiol. 2006;55:805–8.

40. CLSI. Methods for Antimicrobial Dilution and Disk Susceptibility Testing of Infrequently Isolated or Fastidious Bacteria: Approved Guideline. 2010;

41. CLSI. Methods for antimicrobial dilution and disk susceptibility testing of infrequently isolated or fastidious bacteria. 3rd ed. M45 C guideline, editor. Wayne, Penn., USA.: Clinical and Laboratory Standards Institute, CLSI guideline M45; 2016.

42. Dungeni M, van Der Merwe RR, Momba MNB. Abundance of pathogenic bacteria and viral indicators in chlorinated effluents produced by four wastewater treatment plants in the Gauteng Province, South. Water SA. 2010;36:607–14.

43. Ye L, Zhang T. Bacterial communities in different sections of a municipal wastewater treatment plant revealed by 16S rDNA 454 pyrosequencing. Appl. Microbiol. Biotechnol. 2013;97:2681–90.

44. Ngari J, Kotut K, Okemo P. Potential threat to wildlife posed by enteric pathogens from Nakuru sewage treatment plant. Afr. J. Health Sci. 2011;18:85–95.

45. Ramteke PW, Awasthi S, Srinath T, Joseph B. Efficiency assessment of Common Effluent Treatment Plant (CETP) treating tannery effluents. Environ. Monit. Assess. 2010;169:125–31.

46. Odjadjare EEO, Okoh AI. Prevalence and distribution of Listeria pathogens in the final effluents of a rural wastewater treatment facility in the Eastern Cape Province of South Africa. World J. Microbiol. Biotechnol. 2010;26:297–307.

47. Odjadjare E. Prevalence of Listeria pathogens in effluents of some wastewater treatment facilities in the Eastern Cape Province of South Africa. University of Fort Hare; 2010.

48. DEFRA. Review of Operational and Experimental Techniques for the Removal of Bacteria, Viruses and Pathogens from Sewage Effluents. London; 1988.

49. Shang C, Qi Y, Lo IMC. Factors Affecting Inactivation Behavior in the Monochloramination Range. J. Environ. Eng. American Society of Civil Engineers; 2005;131:119–29.

50. Sibanda T, Chigor VN, Okoh AI. Seasonal and spatio-temporal distribution of faecal-indicator bacteria in Tyume River in the Eastern Cape Province, South Africa. Environ. Monit. Assess. 2013;185:6579–90.

51. Henning W. Green Drop [Internet]. 2010 [cited 2014 Jun 17]. Available from: http://www.erwat.com/page.php?pageID=7

52. DWAF: Department of Water Affairs. 2011 Green Drop Report. 2011.

53. DWAF D of WA. Chapter 5 – Eastern Cape Province. Green Drop Rep. 2012. p. 102–49.

54. DWAF: Department of Water Affairs. 2013 Green Drop Report. 2016.

55. DWAF: Department of Water Affairs. 2014 Green Drop Report. 2016.

56. Wong H, You W, Chen S. Detection of Toxigenic Vibrio cholerae, V. pamhaemolyticus and V. vulnificus in Oyster by Multiplex-PCR with Internal Amplification Control. J. Food Drug Anal. 2012;20:48– 58.

57. Drake SL. The Ecology Vibrio vulnificus and Vibrio parahaemolyticus from Oyster Harvest Sites in the Gulf of Mexico. 2008.

58. Vaseeharan B, Ramasamy P, Murugan T, Chen JC. In vitro susceptibility of antibiotics against Vibrio spp. and Aeromonas spp. isolated from Penaeus monodon hatcheries and ponds. Int. J. Antimicrob. Agents. 2005;26:285–91.

59. Lee SW, Najiah M, Wendy W, Nadirah M. Comparative study on antibiogram of Vibrio spp. isolated from diseased postlarval and marketable-sized white leg shrimp (Litopenaeus vannamei). Front. Agric. China. 2009;3:446–51.

60. CDC C for DC and P. Vibrio Illness (Vibriosis) [Internet]. 2013 [cited 2014 Oct 2]. Available from: http://www.cdc.gov/vibrio/

61. Han F, Walker RD, Janes ME, Prinyawiwatkul W, Ge B. Antimicrobial susceptibilities of Vibrio parahaemolyticus and Vibrio vulnificus isolates from Louisiana Gulf and retail raw oysters. Appl. Environ. Microbiol. 2007;73:7096–8.

62. Shaw KS, Rosenberg Goldstein RE, He X, Jacobs JM, Crump BC, Sapkota AR. Antimicrobial susceptibility of Vibrio vulnificus and Vibrio parahaemolyticus recovered from recreational and commercial areas of Chesapeake Bay and Maryland Coastal Bays. PLoS One. Public Library of Science; 2014;9:e89616.

63. Ismail H, Smith AM, Tau NP, Sooka A, Keddy KH. Cholera outbreak in South Africa, 2008-2009: laboratory analysis of Vibrio cholerae O1 strains. J. Infect. Dis. 2013;208 Suppl:S39–45.

64. Obi CL, Bessong PO, Momba MNB, Potgieter N, Samie A, Igumbor EO. Profiles of antibiotic susceptibilities of bacterial isolates and physico-chemical quality of water supply in rural Venda communities, South Africa. 2004;30:515–20.

65. Urassa W, Mhando Y. Antimicrobial susceptibility pattern of Vibrio cholerae 01 strains during two cholera outbreaks in Dar Es Salaam, Tanzania. East African Med 2000;77:350–3.

66. Okuda J, Ramamurthy T, Yamasaki S. Antibacterial activity of ciprofloxacin against clinical strains of Vibrio cholerae O139 recently isolated from India. Yakugaku Zasshi. 2007;127:903–4.

67. Tran HD, Alam M, Trung NV, Kinh N Van, Nguyen HH, Pham VC, et al. Multi-drug resistant Vibrio cholerae O1 variant El Tor isolated in northern Vietnam between 2007 and 2010. J. Med. Microbiol. 2012;61:431–7.

68. Benedicta OE. Antibiotic Susceptibility Patterns and Plasmid Profile of Vibrio cholerae from Water Samples in Elele Community, Rivers State, Nigeria Vibro cholerae was found to concentrate on the the biotype eltor whereas in the Southern coastal regions the 0139 Vibr. J. Appl. Sci. Environ. Manag. March. 2012;16:115–20.

69. Laganà P, Caruso G, Minutoli E, Zaccone R, Santi D. Susceptibility to antibiotics of Vibrio spp. and Photobacterium damsela ssp. piscicida strains isolated from Italian aquaculture farms. New Microbiol. 2011;34:53–63.

70. Morris JG, Sztein MB, Rice EW, Nataro JP, Losonsky GA, Panigrahi P, et al. Vibrio cholerae O1 can assume a chlorine-resistant rugose survival form that is virulent for humans. J. Infect. Dis. 1996;174:1364–8.

71. Quilici ML, Massenet D, Gake B, Bwalki B, Olson DM. Vibrio cholerae O1 variant with reduced susceptibility to ciprofloxacin, Western Africa. Emerg. Infect. Dis. Centers for Disease Control and Prevention; 2010. p. 1804–5.

72. Roychowdhury A, Pan A, Dutta D, Mukhopadhyay AK, Ramamurthy T, Nandy RK, et al. Emergence of tetracycline-resistant Vibrio cholerae O1 Serotype Inaba, in Kolkata, India. Jpn. J. Infect. Dis. 2008;61:128–9.

73. Rashed SM, Hasan NA, Alam M, Sadique A, Sultana M, Hoq MM, et al. Vibrio cholerae O1 with reduced susceptibility to ciprofloxacin and azithromycin isolated from a rural coastal area of Bangladesh. Front. Microbiol. 2017;8.

74. Leclercq R, Cantón R, Brown DFJ, Giske CG, Heisig P, MacGowan a P, et al. EUCAST expert rules in antimicrobial susceptibility testing. Clin. Microbiol. Infect. 2013;19:141–60.

75. Kim H Bin, Wang M, Ahmed S, Park CH, LaRocque RC, Faruque ASG, et al. Transferable quinolone resistance in Vibrio cholerae. Antimicrob. Agents Chemother. 2010;54:799–803.

76. Dalsgaard a, Forslund a, Sandvang D, Arntzen L, Keddy K. Vibrio cholerae O1 outbreak isolates in Mozambique and South Africa in 1998 are multiple-drug resistant, contain the SXT element and the aadA2 gene located on class 1 integrons. J. Antimicrob. Chemother. 2001;48:827–38.

77. Ismail H, Smith AM, Sooka A, Keddy KH. Genetic characterization of multidrug-resistant, extended-spectrum-β-lactamase-producing Vibrio cholerae O1 outbreak strains, Mpumalanga, South Africa, 2008. J. Clin. Microbiol. 2011;49:2976–9.

78. Singh D V, Choudhury R, Panda S. Emergence and dissemination of antibiotic resistance: A global problem. Indian J. Med. Microbiol. 2014;30:384–90.

79. Mandal J, Dinoop KP, Parija SC. Increasing antimicrobial resistance of Vibrio cholerae OI biotype E1 tor strains isolated in a tertiary-care centre in India. J. Health. Popul. Nutr. 2012;30:12–6.

80. Baron S, Larvor E, Chevalier S, Jouy E, Kempf I, Granier SA, et al. Antimicrobial susceptibility among urban wastewater and wild shellfish isolates of Non-O1/Non-O139 Vibrio cholerae from La Rance estuary (Brittany, France). Front. Microbiol. 2017;8.

81. Raissy M, Moumeni M, Ansari M, Rahimi E. Antibiotic resistance pattern of some Vibrio strains isolated from seafood. Iran. J. Fish 2012;

82. CLSI. Performance Standards for Antimicrobial Susceptibility Testing; Twenty-Second Informational Supplement. Clinical and Laboratory Standards Institute; 2012.

83. CLSI. M45. Methods for Antimicrobial Dilution and Disk Susceptibility Testing of Infrequently Isolated or Fastidious Bacterial]; Proposed Guideline. Guidel. CLSI. 2015.

84. Faruque SM, Albert MJ, Mekalanos JJ. Epidemiology, genetics, and ecology of toxigenic Vibrio cholerae. Microbiol. Mol. Biol. Rev. 1998;62:1301–14.

85. Rafi S, Qureshi AH, Saeed W, Ali A, Ahmadani M, Khawaja SA. Changing epidemiology and sensitivity pattern of Vibrio cholerae at Rawalpindi. Pakistan J. Med. Sci. 2004;20:357–60.

86. Marin M a., Thompson CC, Freitas FS, Fonseca EL, Aboderin a. O, Zailani SB, et al. Cholera Outbreaks in Nigeria Are Associated with Multidrug Resistant Atypical El Tor and Non-O1/Non-O139 Vibrio cholerae. PLoS Negl. Trop. Dis. 2013;7.

87. Mandal J, Preethi V, Vasanthraja R, Srinivasan S, Parija SC. Resistance to ceftriaxone in Vibrio cholerae. Indian J. Med. Res. 2012;136:674–5.

88. Wong MHY, Liu M, Wan HY, Chen S. Characterization of extended-spectrum-β-lactamase-producing Vibrio parahaemolyticus. Antimicrob. Agents Chemother. 2012;56:4026–8.

89. Liang P, Cui X, Du X, Kan B, Liang W. The virulence phenotypes and molecular epidemiological characteristics of Vibrio fluvialis in China. Gut Pathog. 2013;5:6.

90. Zanetti S, Spanu T, Deriu A, Romano L, Sechi LA, Fadda G. In vitro susceptibility of Vibrio spp. isolated from the environment. Int. J. Antimicrob. Agents. 2001;17:407–9.

91. Chiang S-R, Chuang Y-C. Vibrio vulnificus infection: clinical manifestations, pathogenesis, and antimicrobial therapy. J. Microbiol. Immunol. Infect. 2003;36:81–8.

92. Larsson DGJ, de Pedro C, Paxeus N. Effluent from drug manufactures contains extremely high levels of pharmaceuticals. J. Hazard. Mater. 2007;148:751–5.

93. Li D, Yang M, Hu J, Zhang Y, Chang H, Jin F. Determination of penicillin G and its degradation products in a penicillin production wastewater treatment plant and the receiving river. Water Res. 2008;42:307–17.

94. Hendricks R, Pool EJ. The effectiveness of sewage treatment processes to remove faecal pathogens and antibiotic residues. J. Environ. Sci. Health. A. Tox. Hazard. Subst. Environ. Eng. 2012;47:289–97.

95. Kümmerer K. Antibiotics in the aquatic environment--a review--part II. Chemosphere. Elsevier Ltd; 2009;75:435–41.

96. Opal SM, Pop-vicas A. Molecular Mechanisms of Antibiotic Resistance in Bacteria. Mand. Douglas, Bennett’s Princ. Pract. Infect. Dis. 8th ed. Elsevier; 2015. p. 235–51.

97. Luo Y, Yang F, Mathieu J, Mao D, Wang Q, Alvarez PJJ. Proliferation of Multidrug-Resistant New Delhi Metallo-β-lactamase Genes in Municipal Wastewater Treatment Plants in Northern China. 2014;1–5.

98. Ohlsen K, Ternes T, Werner G, Wallner U, Löffler D, Ziebuhr W, et al. Brief report Impact of antibiotics on conjugational resistance gene transfer in Staphylococcus aureus in sewage. 2003;5:711–6.

99. Kümmerer K. The presence of pharmaceuticals in the environment due to human use--present knowledge and future challenges. J. Environ. Manage. 2009;90:2354–66.

100. Rizzo L, Manaia C, Merlin C, Schwartz T, Dagot C, Ploy MC, et al. Urban wastewater treatment plants as hotspots for antibiotic resistant bacteria and genes spread into the environment: a review. Sci. Total Environ. Elsevier B.V.; 2013;447:345–60.

